# Same-day diagnostic and surveillance data for tuberculosis via whole genome sequencing of direct respiratory samples

**DOI:** 10.1101/094789

**Authors:** Antonina A. Votintseva, Phelim Bradley, Louise Pankhurst, Carlos del Ojo Elias, Matthew Loose, Kayzad Nilgiriwala, Anirvan Chatterjee, E. Grace Smith, Nicholas Sanderson, Timothy M. Walker, Marcus R. Morgan, David H. Wyllie, A. Sarah Walker, Tim E. A. Peto, Derrick W. Crook, Zamin Iqbal

## Abstract

Routine full characterization of *Mycobacterium tuberculosis* (TB) is culture-based, taking many weeks. Whole-genome sequencing (WGS) can generate antibiotic susceptibility profiles to inform treatment, augmented with strain information for global surveillance; such data could be transformative if provided at or near point of care.

We demonstrate a low-cost DNA extraction method for TB WGS direct from patient samples. We initially evaluated the method using the Illumina MiSeq sequencer (40 smear-positive respiratory samples, obtained after routine clinical testing, and 27 matched liquid cultures). *M. tuberculosis* was identified in all 39 samples from which DNA was successfully extracted. Sufficient data for antibiotic susceptibility prediction was obtained from 24 (62%) samples; all results were concordant with reference laboratory phenotypes. Phylogenetic placement was concordant between direct and cultured samples. Using an Illumina MiSeq/MiniSeq the workflow from patient sample to results can be completed in 44/16 hours at a cost of £96/£198 per sample.

We then employed a non-specific PCR-based library preparation method for sequencing on an Oxford Nanopore Technologies MinION sequencer. We applied this to cultured *Mycobacterium bovis* BCG strain (BCG), and to combined culture-negative sputum DNA and BCG DNA. For the latest flowcell, the estimated turnaround time from patient to identification of BCG was 6 hours, with full susceptibility and surveillance results 2 hours later. Antibiotic susceptibility predictions were fully concordant. A critical advantage of the MinION is the ability to continue sequencing until sufficient coverage is obtained, providing a potential solution to the problem of variable amounts of *M. tuberculosis* in direct samples.

## Introduction

The long-standing gold standard for *Mycobacterium tuberculosis* drug susceptibility testing (DST) is the phenotypic culture-based approach, which is time-consuming and laborious. First-line tuberculosis (TB) treatment includes four drugs (rifampicin, isoniazid, ethambutol and pyrazinamide) but with the spread of multi-drug resistant strains, there is a growing need for data on second-line drugs, including the fluoroquinolones, and aminoglycosides.

Due to long turnaround times for phenotypic testing (up to two months), these are often preceded by WHO-endorsed molecular methods such as the GenoType MTBDRplus and MTBDRsl assays (Hain Lifescience GmbH, Germany), and Xpert MTB/RIF (Cepheid, USA). These potentially culture-free, PCR-based tests rapidly identify species and detect the most common drug resistance conferring mutations. However, this technology is limited by the number of mutations that can be probed. This limitation is of concern, given the many low frequency drug resistance conferring mutations in *M. tuberculosis*, particularly for second-line drugs (1). Consistent with this concern, the proportion of phenotypically resistant samples which are detectable by MTBDRplus range from 21-25% for the second-line drugs capreomycin and kanamycin (2) to 98.4% and 91.4% for the critical first-line drugs rifampicin and isoniazid (3). A potential solution is to sequence amplicons targeting a wider range of resistance conferring genes, as previously demonstrated (4).

The potential of whole genome sequencing (WGS) as a diagnostic assay has been repeatedly identified (5–7). Recent studies based on WGS of mycobacteria have evaluated WGS-based susceptibility predictions (1, 8–10), species identification, and elucidation of epidemiology (11–16). This has culminated in the first successful application of WGS as a clinical diagnostic for mycobacteria from early positive liquid cultures (16). Moreover, WGS was performed at a cost comparable with existing phenotypic assays and offered faster turnaround times.

Generating WGS information directly from patient samples, and avoiding the need for culture, would be transformative. However, direct samples contain highly variable amounts of mycobacterial cells mixed with other bacterial and human cells; the latter accounting for up to 99.9% of DNA present. Furthermore, mycobacterial cells may aggregate due to the high mucus content of some samples; meaning sample volume and Acid Fast Bacillus (AFB) count may not represent the total quantity of mycobacteria available. Direct samples therefore require pre-processing to homogenize and enrich for mycobacteria by depleting other cells/DNA. The challenges of direct sample processing were illustrated by two studies assessing the feasibility of WGS directly from clinical samples (17–18). By sequencing eight smear-positive sputum samples subjected to differential lysis followed by DNA extraction with a commercial kit, Doughty and colleagues were able to achieve only 0.002-0.7X depth of coverage for *M. tuberculosis* with 20.3-99.3% of sequences mapping to the human genome. 7/8 samples could be assigned to *M. tuberculosis* complex, but none had sufficient data for drug susceptibility prediction. In a second study, Brown and colleagues applied a SureSelect target enrichment method (Agilent, USA) to capture *M. tuberculosis* DNA prior to WGS. 20/24 smear-positive samples achieved 90% genome coverage with ≥20xdepth; sufficient for prediction of species and antibiotic susceptibility. However, the protocol was slow (2-3 days) and may be prohibitively expensive for use in low-income settings.

In this study we test a simple low-cost DNA extraction method using Illumina MiSeq WGS on 40 smear-positive, primary respiratory samples from *M. tuberculosis* infected patients. We evaluate the protocol in terms of DNA obtained, species assignment of the sequenced reads, and our ability to obtain key clinical data (detection of *M. tuberculosis* and antibiotic susceptibility prediction) along with epidemiological information (placement on phylogenetic tree). These data would enable a single test to deliver the core information for both patient and public health in <48 hours using Illumina-based WGS. We also develop an approach for WGS using the highly portable, random-access, Oxford Nanopore Technologies (ONT) MinION, reducing potential turnaround time to 8 hours.

## Materials and Methods

### Sample selection and processing

Direct respiratory Ziehl-Neelsen (ZN)-positive samples with acid-fast bacilli (AFB) scorings from +1 to +3 were collected from patients with subsequently confirmed *M. tuberculosis* infections at the John Radcliffe Hospital, Oxford Universities NHS Foundation Trust, Oxford, UK (n=18), and Birmingham Heartlands Hospital NHS Foundation Trust, Birmingham, UK(n=22). 2/18 Oxford samples were culture negative specimens taken 2.5 months apart from the same patient undergoing treatment for *M. tuberculosis.* If available, corresponding Mycobacterial Growth Indicator Tube (MGIT) cultures were collected for each direct sample (Oxford n=11, Birmingham n=17). Two ZN and culture negative direct respiratory samples were also collected from the John Radcliffe Hospital.

The discarded direct samples were collected only after sufficient material had been obtained for the routine diagnostic workflow, including the requirement to ensure that enough sample volume remained if re-culture was requested. Consequently, our samples were of lower volume and quality than would be the case if the method were used routinely. While waiting for the routine laboratory results, samples were stored at +4C and later processed in batches of 5-12. All ZN-positive samples were digested and decontaminated with NAC-PAC RED kit (AlphaTec, USA). Direct samples and corresponding MGIT culture aliquots (1 mL) were heat inactivated in a thermal block after sonication (20 min, 35 kHz) for 30 min and 2 h at 95C, respectively. MGITs were inactivated for 2 hours owing to their high bacterial load. Before DNA extraction samples were stored at +4C.

### DNA extraction and Illumina MiSeq sequencing

Mycobacterial DNA from MGIT cultures was extracted using a previously validated ethanol precipitation method (19). DNA from ZN-positive direct samples was extracted using a modified version of this protocol. These modifications included a saline wash followed by MolYsis Basic5 kit (Molzym, Germany) treatment for the removal of human DNA, and addition of GlycoBlue co-precipitant (LifeTechnologies, USA) to the ethanol precipitation step (Supplementary Figure 1).

Libraries were prepared for the MiSeq Illumina sequencing using a modified Illumina Nextera XT protocol (19). Samples were sequenced using the MiSeq Reagent Kit v2, 2 x 150bp in batches of 9-12 per flow-cell.

### DNA extraction for ONT MinION and Illumina MiniSeq sequencing

ZN/culture-negative sputum and BCG (Pasteur strain; cultivated at 37C in MGIT tubes) DNA was extracted using a modified version of that in (19). Briefly, following a saline wash, samples were re-suspended in 100 μL of molecular grade water and subjected to three rounds of bead-beating at 6 m/s for 40 seconds. The beads were pelleted by centrifugation at 16,100 xg for 10 minutes and 50 μL supernatant cleaned using 1.8x volume AMPure beads (Beckman Coulter, UK). Samples were eluted in 25 μL molecular grade water, and quantified using the Qubit fluorimeter (Thermo Fisher Scientific, USA). (Steps I, III, V and VI of Miseq protocol, Supplementary Figure 1.)

### MiniSeq sequencing

Extracted ZN-negative sputum DNA and pure BCG DNA were combined in a 50:50 ratio (0.5 ng each) and libraries prepared alongside pure BCG DNA (1 ng) using a modified Illumina Nextera XT protocol (19). BCG and two BCG+sputum DNA samples were sequenced at Illumina Cambridge Ltd. UK, using a Mid Output kit (FC-420-1004) reading 15 tiles and with 101 cycles.

### MinION sequencing

All MinION sequencing utilized the best sample preparation kits and flow cells available at the time. A single ZN-negative sputum extract was divided into three equal concentration aliquots (187 ng), and BCG DNA added at 5%, 10% and 15% of the total sputum DNA concentration. These 5-15% spikes represent the lower end of the spectrum seen in the MiSeq samples above (see Figure 2a). These samples, along with pure BCG DNA, were prepared following ONTs PCR-based protocol for low-input libraries (DP006_revB_14Aug2015), using modified primers supplied by ONT, a 20 ng DNA input into the PCR reaction, and LongAmp *Taq* 2X Master Mix (New England Biolabs, USA). PCR conditions were as follows: initial denaturation at 95°C for 3 minutes, followed by 18 cycles of 95°C for 15s, 62°C for 15s, and 65°C for 2.5 minutes, and a final extension at 65°C for 5 minutes. Samples were cleaned in 0.4x volume AMPure beads and the PCR product assessed using the Qubit fluorimeter and TapeStation (Agilent, UK). The final elution was into 10 μL 50 mM NaCl, 10 mM Tris.HCl pH8.0. Finally, 1 μL of PCR-Rapid Adapter (PCR-RAD; supplied by ONT) was added and samples incubated for 5 minutes at room temperature to generate pre-sequencing mix. The pre-sequencing mix was prepared for loading onto flow cells following standard ONT protocols, with a loading concentration of 50 – 100 fmol.

Using the 15% BCG spiked sputum DNA prepared above, amplification was repeated using Phusion High-Fidelity PCR Master Mix with DMSO (New England BioLabs, USA). Gradient PCR was performed to identify the optimal annealing temperature for recovery of BCG DNA (data not shown). Final PCR conditions were as follows: initial denaturation at 98°C for 30s, followed by 18 cycles of 98°C for 10s, 59°C for 15s, and 72°C for 1.5 minutes, and a final extension of 72°C for 10 minutes. Following PCR, the sample was prepared for sequencing as described above. The final loading concentration was approximately 27 fmol.

The above samples were sequenced using R9 spot-on generation flow cells and the 48-hour protocol for FLO-MIN105 (ONT, UK). Base calling was performed via the Metrichor EPI2ME service (ONT, UK) using the 1D RNN for SQK-RAD001 v1.107 workflow.

Subsequently, a new 15% BCG spiked sputum was prepared as described above using Phusion Master Mix with DMSO. Sequencing was performed using R9.4 spot-on generation flow cells and the 48-hour FLO-MIN106 protocol (ONT, UK). Final loading concentration was 43 fmol. Base calling was performed after sequencing was complete using Albacore (ONT, UK), as base calling via Metrichor failed.

### Bioinformatic analysis of Illumina data

To determine levels of contamination and *M. tuberculosis* in samples, reads were immediately mapped using bwa_mem (20) to the human reference genome GRCh37 (hg19) and human reads counted and permanently discarded. Remaining stored reads where then mapped to the *M. tuberculosis* H37Rv reference strain (GenBank NC_018143.2), and any unmapped reads were then mapped to nasal, oral and mouth flora available in the NIH Human Microbiome Project database (http://www.hmpdacc.org/). A minimum reference genome coverage depth of 5 was required for phylogenetic analysis to proceed.

Mycobacterial species and antibiotic resistance to isoniazid, rifampicin, ethambutol, pyrazinamide, streptomycin, aminoglycosides (including capreomycin, amikacin and kanamycin) and fluoroquinolones (including moxifloxacin, ofloxacin, and ciprofloxacin) was predicted using Mykrobe predictor software (21) v0.3.5 updated with a new validated catalogue of resistance conferring genetic mutations (1). For samples where the estimated depth of kmer-coverage of *M. tuberculosis* reported by Mykrobe predictor fell below 3x, no resistance predictions were made. The precise command used was:, “mykrobe predict SAMPLE_ID tb −1 FASTQ -panel walker-2015 -min-depth 3”.

### Phylogenetic analysis

Conservative SNP calls were made using Cortex (22) (independent workflow, k=31) on 3480 samples from (1). Singleton variants were discarded, and a de-duplicated list of 68695 SNPs was constructed. All samples (from our study and from (1)) were genotyped at these sites using the Cortex genotyping model (22). We then measured the number of SNP differences between paired direct and MGIT samples, counting only sites where both genotypes had high confidence (difference between log likelihood of called genotype and of uncalled genotype greater than 1), and neither site was called as heterozygous.

Samples were placed on the phylogenetic tree of 3480 samples from (1) by identifying the leaf with the fewest SNP differences, across the 68695 sites. Placement therefore returns a closest leaf, and a SNP distance.

### Statistical analysis

Univariable and multivariable linear regression was used to identify independent factors affecting log10 DNA concentration after extraction. Analyses were performed using Stata 14.1 (2015, StataCorp, USA).

### Bioinformatic Analysis of MinION Data

Mykrobe predictor version v0.3.5-0-gd724461 was used to predict resistance from the MinION basecalled reads (command: mykrobe predict SAMPLE_ID tb −1 FASTQ -panel walker-2015 - min-depth 3).

As for Illumina data, kmer coverage was estimated using a set of species-informative sequence probes defined in (21), and susceptibility predictions were only made if the median coverage was >3. Yield and timing was analysed using Poretools (23). For the R9.4 sample, Mykrobe predictor was applied to the cumulative read output at each hour. We excluded one false positive rifampicin resistance call seen at low coverage (the first 5 hours of sequencing) that was due to a known software bug (see Supplementary Text 1). Yield of BCG was measured by mapping to a BCG reference (accession BX248333.1).

Phylogenetic placement of the 15% spike BCG sample sequenced on MinION R9.4 was achieved as for the Illumina data - by genotyping 68695 SNPs using the Cortex model, and choosing the leaf with the fewest SNP differences across those sites.

### MinION error analysis

Error bias in the consensus of MinION R9 1D pure BCG reads was measured in two ways, using reads from the pure BCG sequencing run mentioned above.

1. Reads were mapped to the *M. tuberculosis* reference genome using bwa_mem, and then this was passed to the consensus tool racon (24). The output of this was compared with the BCG reference genome using MUMMER (25). Any observed SNPs were considered to be errors, and bias in these errors was observed by looking at isolated SNPs (avoiding alignment artefacts due to nearby indels). The results are shown in Supplementary Table 1.
2. A *de novo* assembly was performed with Canu (26), and then this was compared with the BCG reference genome using MUMMER, as above. Full results shown in Supplementary Table 2.

The results were broadly consistent between the two methods. The mapping approach (number 1 above) found 28% of consensus errors were A->G and 60% were T->C. (Note these refer to the SNP with respect to the reference, not to errors within a single read.) The de novo assembly approach found 50% of consensus errors were A->G, and 44% were T->C.

### Costing analysis

Basic costing included reagents required for sample decontamination, DNA extraction, MiSeq and Nanopore library preparations, and sequencing; correct as of November 2016. Generic laboratory consumables (e.g. pipette tips, tubes) were not included. SureSelect (Agilent, UK) costs, as used by Brown *et al.* (18), were obtained via a company representative and were correct of June 2016. United States Dollars (USD) were converted to Great British Pounds (GBP) at $1.25 USD per GBP. See Supplementary Table 3 for details.

### Ethics

For this study no ethical review was required because it was a laboratory methods development study focusing on bacterial DNA extracted from discarded samples identified only by laboratory numbers with no personal or clinical data. Sequencing reads identified as human based on fast mapping with BWA were counted and immediately permanently discarded (i.e. never stored electronically).

### Accession numbers

Genome sequence data are in the process of being deposited to the Sequence Read Archive (SRA), NCBI. The MiSeq data is submitted under the study accession number SRP093599. The MiniSeq and MinION data accession numbers will be entered here when available.

## Results

### DNA extraction protocol and evaluation of Illumina sequencing output

DNA was extracted from 40 ZN-positive direct respiratory samples, of which 38 were culture-confirmed *M. tuberculosis* (“culture-positive”) and 2 were culturenegative. DNA was also extracted from 28 available corresponding MGIT cultures. All direct samples were the remainder of specimens available after processing by the routine laboratory, and therefore had variable volume (median 1.5 ml, IQR 0.5-3.1, range 0.25-15) and age (median 30 days from collection to processing, IQR 15-45, range 0-67). Most direct samples (78%; 31/40) could therefore be considered suboptimal on the basis of either low volume (≤1ml) or long storage time (≥30 days) or both.

After DNA extraction, 33/40 (83%) direct samples and all 28 MGIT cultures yielded ≥0.2 ng/μl DNA, the amount recommended for MiSeq Illumina library preparation (Figure 1).

**Figure 1:**
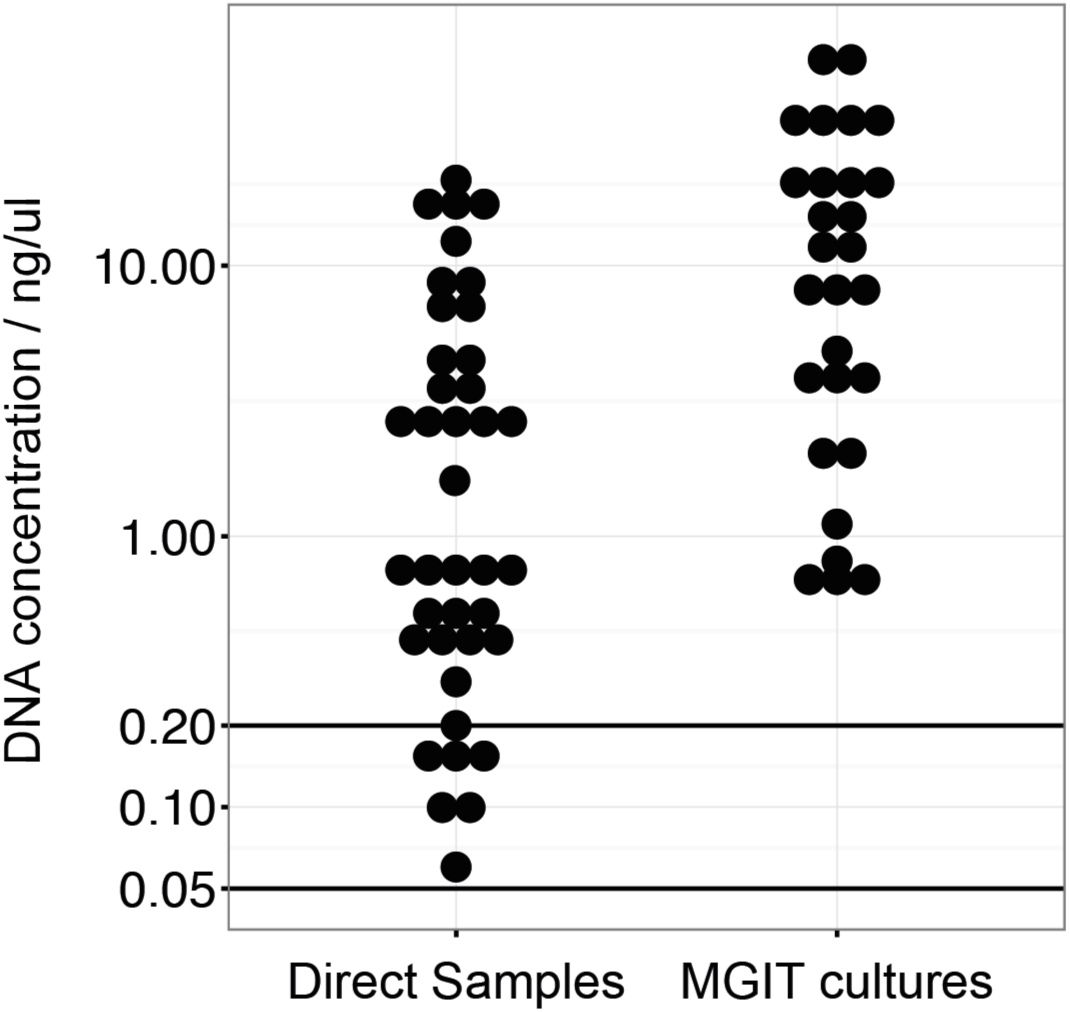
DNA extracted (ng/μl) from MGIT cultures and direct clinical samples. Each dot represents a single extraction. Horizontal line at 0.2 ng/μl represents the DNA concentration required for MiSeq library preparation. Horizontal line at 0.05 ng/μl represents minimum DNA concentration used for MiSeq library preparation from direct samples.

There was no evidence that DNA yield was affected (either in multivariable or univariable models) by (1) sample type (sputum or bronchoalveolar lavage) (p=0.94; univariable linear regression), (2) AFB scorings (from +1 to +3) (p=0.37), (3) storage time prior to DNA extraction (days from collection) (p=0.51) and (4) sample volume (p=0.28). Although DNA concentration was measured and recorded after extraction, further data on DNA quality (e.g. DNA Integrity Number (DIN) provided by TapeStation (Agilent, USA)) were not routinely recorded.

In total 39/40 direct samples with detectable DNA (37 culture-positive, 2 culture-negative) and 27/28 MGIT cultures were sequenced on an Illumina MiSeq. One MGIT culture was not sequenced because the corresponding direct sample failed to yield measurable DNA. We used a lower than recommended DNA concentration threshold for MiSeq library preparation (>0.05 ng/μl rather than >0.2 ng/μl) on the basis of previous experience of sequencing mycobacterial cultures with suboptimal amounts of DNA (19). 6/40 (15%) samples yielded DNA below the 0.2 ng/μl threshold. All sequenced direct samples produced ≥1.5 million reads (median 3.6 million, IQR 2.9-5.0, range 1.5-12), as did all MGIT cultures (median 3.1 million, IQR 2.8-3.3, range 2.0-4.1).

### Contamination levels of direct and MGIT samples

We assigned reads to categories *M. tuberculosis*, human, naso-pharyngeal flora (NPF) and “other” by mapping (see Methods). 77% (30/39) of direct samples contained <10% human reads. However, only 46% (18/39) contained <10% NPF and other bacterial reads, and 26% (10/39) contained >40% of reads from non-mycobacterial, non-NPF, bacteria (Figure 2a). By comparison, MGIT culture samples showed much less contamination, as expected. (Figur2be 2b).

**Figure 2:**
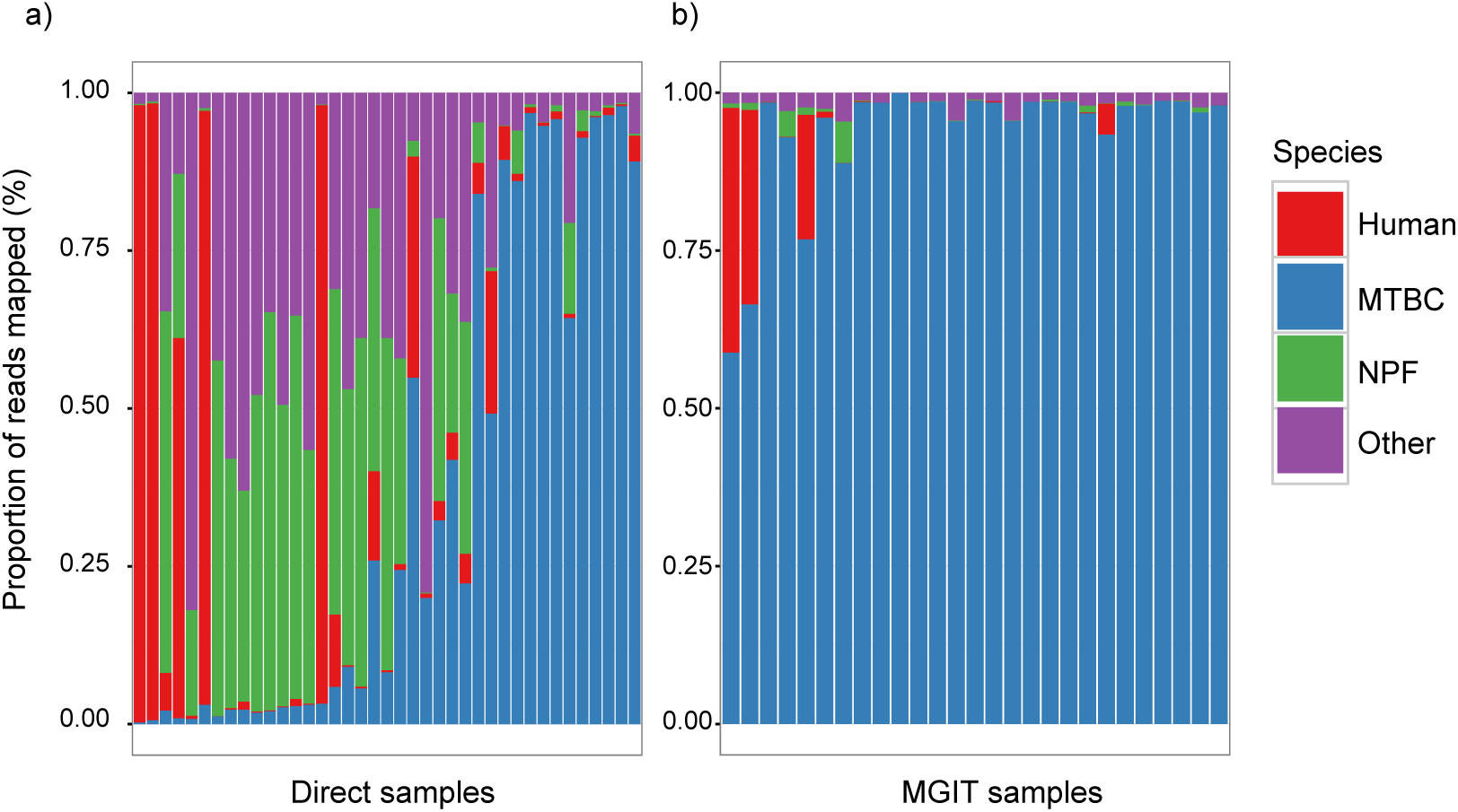
Proportion of reads assigned to various species categories in each sample. **a**) Direct samples show removal of human DNA (red) has been broadly successful, but removal of naso-pharyngeal flora (NPF, green) and other bacteria (purple) had more variable success. **b**) MGIT samples show much more uniform dominance of *M. tuberculosis* reads, as expected after 2 weeks of culture designed to favour mycobacterial growth.

### Recovery of *M. tuberculosis* genome

Figure 3a shows the distribution of the *M. tuberculosis* reference genome depth of coverage across direct samples. Samples either have more than 10x depth and recover more than 90% of the genome, or have <3x depth and recover less than 12% of the genome. The vertical dotted line delineates our threshold of 3x coverage, below which resistance predictions were not made. Figure 3b shows the amount of contamination (reads not mapping to *M. tuberculosis)* per sample. Ten samples had <15% contaminant reads, although contamination levels increased as high as 75% before the proportion of the *M. tuberculosis* genome recovered started to drop. Low numbers of *M. tuberculosis* reads could also reflect poor DNA quality from samples stored for long periods, as most of the samples with <80% reference genome coverage (12/17, 71%) were more than 3 weeks old before extraction.

**Figure 3:**
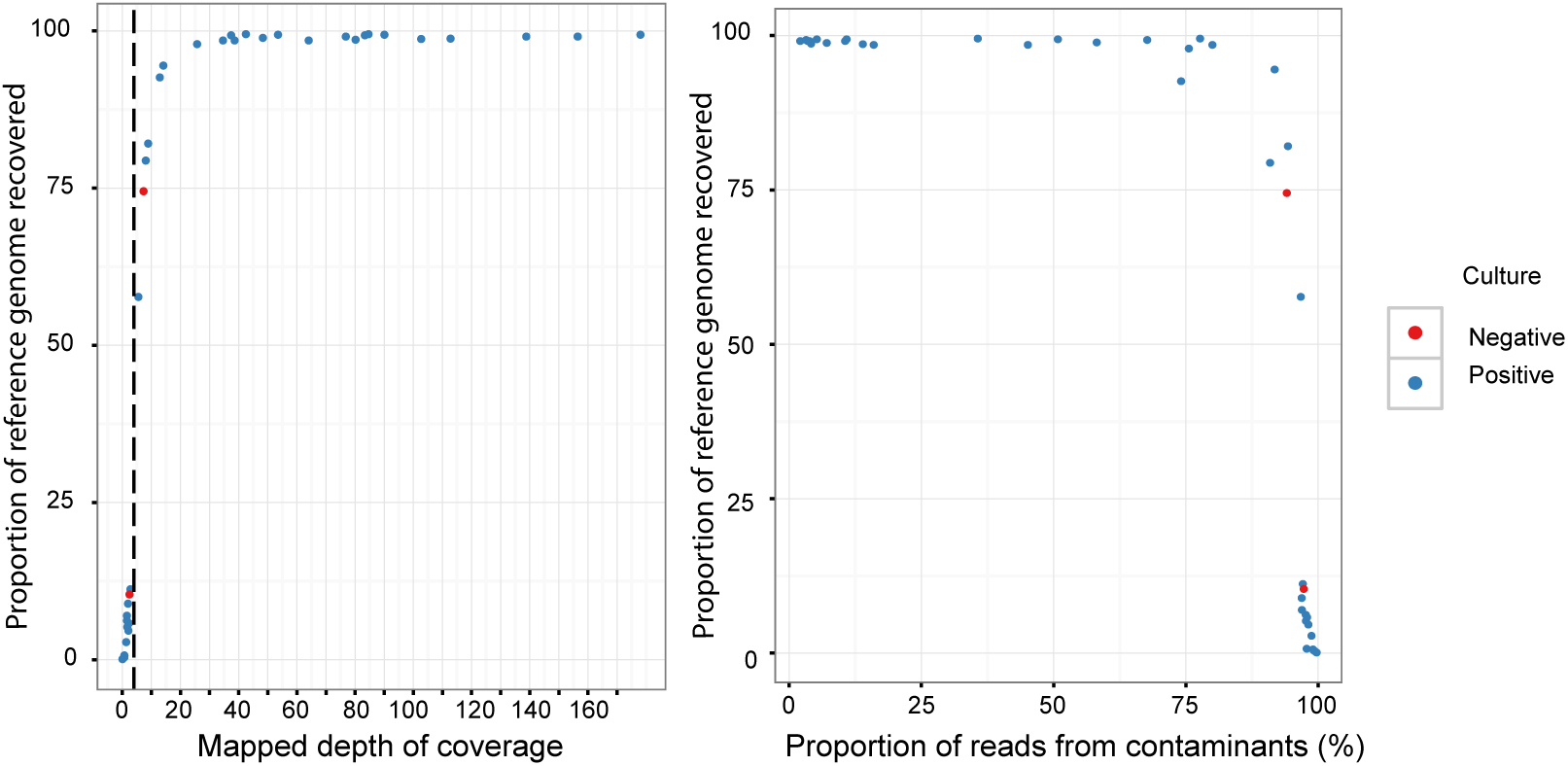
Recovery of *M. tuberculosis* genome in direct samples and robustness to contamination. **a**) Depth versus proportion of the *M. tuberculosis* reference recovered (at >5x depth). Samples either have more than 10x depth and recover more than 90% of the genome, or have <3x depth and recover less than 12% of the genome. Vertical dotted line at 3x depth is threshold for susceptibility prediction. **b**) Proportion of contamination (reads not mapping to *M. tuberculosis* reference) versus proportion of genome recovered. Samples with less than 95% of the *M. tuberculosis* genome recovered all have >75% of contamination.

### Concordance of results from direct and MGIT samples

We took a set of 68,695 high quality SNPs obtained from analysis of 3480 samples (1), and genotyped all samples at these positions (see Methods). This allowed us to calculate a genetic distance between the 17 paired MGIT and direct samples. The median (and modal) SNP difference was 1 (Figure 4a), with one outlier pair of samples that differed by 1106 SNPS, discussed below and all other differences ≤22 SNPs.

**Figure 4:**
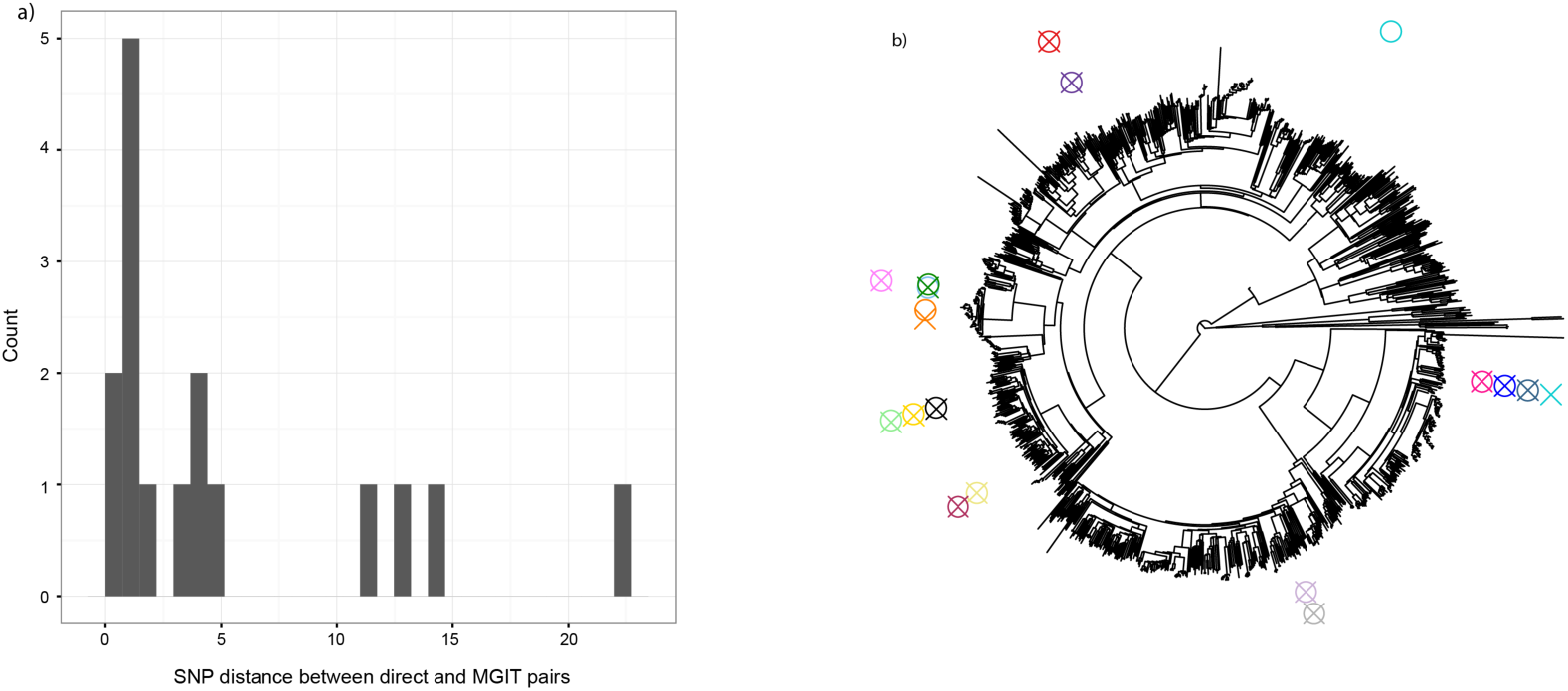
Genotypic concordance between direct and paired MGIT samples. **a**) Histogram of genetic (SNP) differences, excluding the one pair which differ by 1106 SNPs; median (and modal) difference is 1 - thus direct sequencing is identifying the same strain of *M. tuberculosis* as culture-based sequencing would. **b**) Placing direct/MGIT pairs on a phylogenetic tree of 3480 samples shows distribution of samples across world diversity, and for the 1 pair (of 17) with 1106 differences, the MGIT sample places very close to other samples (0 SNP differences to one, 5 SNP differences to others), and so is possibly due to a labeling error.

We placed 17 paired direct and MGIT samples on the phylogeny from (1) (see Methods). Our samples were distributed across global diversity (Figure 4b; tree thinned to aid visibility). For the pair with 1106 SNP differences, the MGIT sample was placed very closely to 3 other pairs (0 SNP difference to one sample, and 5 SNP differences to the others). Although this might result from different strains being present within the host, a within-laboratory labelling error or cross-contamination is also possible.

### No evidence of higher diversity in direct samples

Comparing direct/MGIT pairs where both samples had at least 20x mean depth of coverage on the *M. tuberculosis* reference, the median number of high confidence (see Methods) heterozygous sites was 25 in both direct and MGIT samples. There was no clear trend of greater genome-wide diversity in direct samples (Supplementary Figure 2).

### Detection of *M. tuberculosis* in culture positive/negative samples

All sequenced culture-positive (37/39) direct *M. tuberculosis* samples were successfully identified by Mykrobe predictor to complex level (37/37) and 95% to species level (35/37), including 13/37 (35%) where the mean depth of coverage was <3. All MGIT cultures were identified as *M. tuberculosis.* We were also able to identify *M. tuberculosis* in 2/2 direct samples with low AFB scores (+1) and no growth in MGIT culture; this may represent dead bacilli from a patient undergoing treatment.

### Antibiotic resistance prediction

In total 168 predictions for first-line (n=96) and second-line (n=72) antibiotic resistance were made for the 24/37 (65%) direct samples which had at least 3x depth (Supplementary Tables 4,5). For the 13/37 (35%) samples that had <3x depth, no resistance predictions were made. This included 1/2 culture-negative samples.

92/96 (96%) predictions for the first-line antibiotics were concordant with reference laboratory DST. The four mismatches (three pyrazinamide mixed genotypes with both R and S alleles present, and one rifampicin resistant genotype with sensitive phenotype) were found across three samples, all from the same patient (patient 2 in Supplementary Table 5) who had a variable phenotype for rifampicin and pyrazinamide. The resistant genotype for rifampicin was consistent across all three samples from this patient (rpoB_I491F). There is evidence that this mutation causes resistance, but that the phenotype is often reported as sensitive (27,28,1). The mixed genotype for pyrazinamide was again consistent with presence of both R and S alleles on pncA_V7L across all three samples, whereas the phenotype varied. This mutation is also known to confer resistance in samples reported as phenotypically sensitive (1). Further, 1/3 samples from this patient (sample 602112, Supplementary Table 4) contained two additional mutations conferring resistance to isoniazid and pyrazinamide, katG_S315T and pncA_T135P respectively, which were not detected in the previous or following sample. This variation between same-patient samples taken over time may represent ongoing evolution, changing population size, and within-patient diversity of *M. tuberculosis* as previously demonstrated by Eldholm *et al* (29). In addition to the above, WGS provided 72 predictions for second-line antibiotics where DST was not attempted.

The 13/37 samples that yielded insufficient WGS data for resistance prediction had a higher proportion of other bacterial DNA (Figure 3b; median 96%, IQR 3870%, vs median 12%,IQR 0-67%, in those where resistance prediction was possible, rank-sum p=0.01).

### Sub-24 hour turnaround time with Illumina MiniSeq

Illumina MiniSeq sequencing for three samples (single run; 1 pure BCG, 2 negative sputum DNA spiked with BCG DNA) was completed in 6 hours 40 minutes. BCG reference genome coverage was 31-33x in spiked samples, and 84x in pure BCG (Table 1). In all cases the species/strain was correctly identified as *M. bovis* strain BCG, and pyrazinamide resistance was correctly identified due to mutation H57D in *pncA.*

**Table 1:** Yield from pure BCG, and from negative sputum spiked with BC - sequenced on Illumina MiniSeq.

|  | estimated Fmol loaded | Yield (Mb) | Read length (bp) | BCG reference coverage |
| --- | --- | --- | --- | --- |
| Pure BCG TB1_N716 | 800 | 381 | 101 | 84.0 |
| 50 % BCG TB1_N718 | 800 | 244 | 101 | 31.0 |
| 50% BCG TB1_N719 | 800 | 257 | 101 | 33.0 |

### Modified method for ONT MinION

A new PCR-based rapid 1D MinION protocol was tested using extracted BCG DNA, ZN-negative sputum DNA spiked with BCG DNA, and R9 flowcells (see Methods). Analysis of genome-wide coverage distribution confirmed that use of PCR had not led to significant coverage bias (Supplementary Figure 3), and that >95% of the reference genome attained coverage >5x for all samples. In all cases Mykrobe correctly identified the species/strain as *M. bovis* strain BCG (Table 2). Amplification with Phusion High-Fidelity master mix resulted in the highest yield (760Mb, with 68x coverage of BCG). The pure BCG and both 15% spike experiments resulted in correct identification of the H57D mutation in *pncA* that confers pyrazinamide resistance in BCG.

**Table 2:** Yield from pure BCG, and from negative sputum spiked with BCG - both sequenced with MinION 1D protocol.

| Model | Sample | Fmol loaded | Read count | Yield /Mb | Avg read length/ kb | BCG covg depth | SNPs (kmer covg on R allele) |
| --- | --- | --- | --- | --- | --- | --- | --- |
| R9 | Pure culture d BCG | Not available | 297,239 | 360 | 1.2 | 80 | H57D (13) |
| R9 | 5% BCG LongAmp | 82 | 182,670 | 559 | 2.0 | 19 | No calls |
| R9 | 10% BCG LongAmp | 76 | 180,507 | 467 | 1.8 | 10 | No calls |
| R9 | 15% BCG LongAmp | 51 | 203,285 | 627 | 2.0 | 35 | H57D (3) |
| R9 | 15% BCG Phusion | 27 | 184,895 | 758 | 2.4 | 68 | H57D (8) |
| R9.4 | 15% BCG Phusion | 43 | 754,338 | 1306 | 1.7 | 147 | H57D (20) |

No susceptibility predictions were made in the 5% and 10% spike experiments, owing to low coverage. The19x and 10x coverage depth, respectively in these samples, corresponded to kmer coverages (k=15) <3x due to sequencing errors. Higher coverage depth did permit resistance prediction; for example, the LongAmp/15% sample had 35x coverage depth, corresponding to kmer coverage of 3x at the H57D allele. For each of the 3 samples with enough coverage to allow susceptibility predictions, 200 resistance SNPs/indels were genotyped by Mykrobe and no false resistance calls were made. Using two independent methods (see Methods) we found a strong bias in the distribution of SNP errors in the consensus of the MinION data. Both methods agreed the bias was systematic though not on the precise proportions (e.g. of SNP errors in de novo assembly, 50% were A->G, and 44% were T->C; Supplementary Table 1,2), consistent with a strong A->G error bias within a 1D read. A filter to ensure SNP calls have support from reads mapping to both strands could remove such errors.

In all 5 samples sequenced on R9 flowcells, data yield was highest at the start of the run, with consistent yield profiles. For the Phusion/15% run we obtained over 65% of the data in 8 hours, and 80% in 10 hours (Supplementary Figure 4). Thus we were able to detect BCG and predict the correct resistance with 10 hours of sequencing despite the high error rate (Supplementary Figure 5).

### Sub-12 hour turnaround time with ONT R9.4 MinION

We sequenced a single sample (15% BCG spiked ZN-negative sputum) on the latest R9.4 MinION flowcell (see Methods). Yield was 1.3Gb in 48 hours; we were able to detect *M. tuberculosis* complex and identify the strain as BCG within 1 hour of sequencing, and correctly predict pyrazinamide resistance after 2 further hours. The additional yield would enable 3-5 samples to be sequenced per flow cell. Currently, multiplexing capabilities are limited by contamination, and the kmer-coverage attained on susceptible/resistance alleles.

Although in this instance we performed base-calling (conversion of raw MinION electrical output to DNA reads) after sequencing completed (taking 48 hours on a quad-core desktop computer), we subsequently verified (on other samples) that real-time base-calling can be performed using ONTs MinKNOW software. This would give a turnaround time of 8 hours (Figure 5).

**Figure 5:**
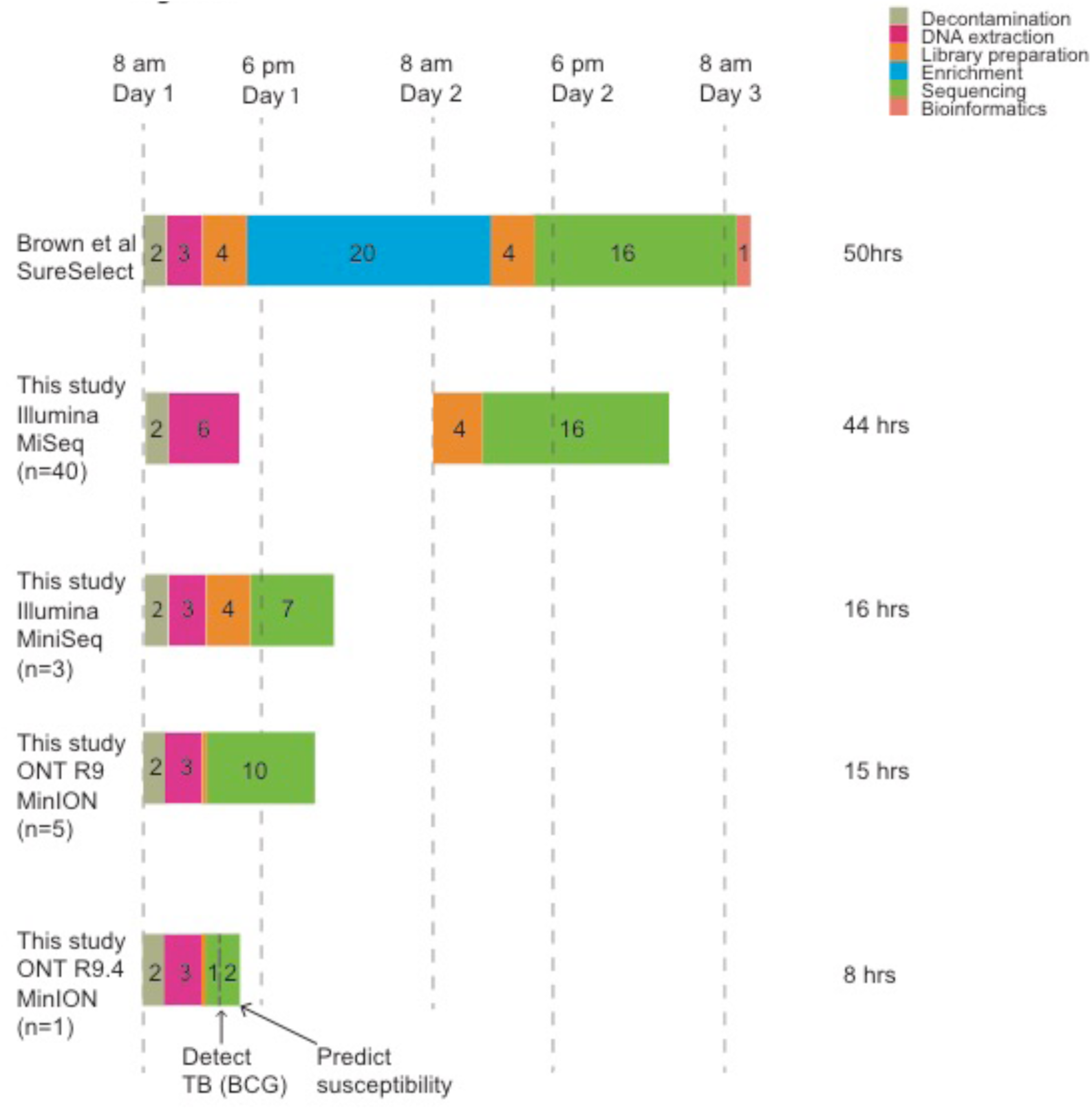
Timelines and cost. We compare the method of Brown et al with the results of this study, using the Illumina MiSeq and MiniSeq, and the ONT MinION. We assume that no step of the process can be initiated after 6pm or before 8am. The method of Brown et al has a rapid extraction step, but also a 20 hour enrichment step, resulting in a 50 hour turn-around time, at an estimated cost of £203/sample. By contrast, our extraction method with MiSeq sequencing provides results at £94/sample. The DNA extraction process was updated for the MiniSeq and MinION experiments, removing the ethanol precipitation step. In normal use this would take 3 hours. The thin orange rectangle on the MinION timelines is the 10 minute sample preparation step. In this experiment, since we used spiked BCG DNA in sputum, we did not use a human depletion step, thus taking only 2 hours. This figure is intended to show comparable real-use timelines, and so the MiniSeq/MinION timelines are shown with 3 hour extraction steps. The MiniSeq enables a 16-hour turnaround time, by sequencing for only 7 hours. The R9 MinION also delivers sub-24 hour results, but requires one flow-cell per sample. The R9.4 MinION gives an 8 hour turnaround time (3 hours of sequencing with real-time (i.e. simultaneous) basecalling when used on a single sample). We do not display the 48 hours we actually took to basecall the data after the run, as this was in effect our error - we subsequently confirmed (on other samples) that real-time basecalling would have worked.

We took our phylogenetic placement of the MiniSeq BCG data as truth, 4 SNPs distant from a BCG sample on the predefined tree. By comparison, after 1 hour of sequencing with R9.4, we were able to genotype 15592 of the 68695 SNPs, placing the sample at the correct leaf of the tree, at an estimated distance of 25 SNPs. Thus, our genotyping on 1D nanopore reads had at most 29 errors out of 15592 SNPs - an error rate of 0.2%.

### Costing

Reagent costs (sample decontamination, extraction, sequencing library preparation, and sequencing) per sample were £96 (MiSeq, 12 samples/run), £198 (MiniSeq, 3 samples/run), £515 (R9 MinION, 1 sample/run), £101-172 (R9.4 MinION, between 3 and 5 samples/run, approximate cost as multiplexing kit not yet available). See Supplementary Table 3 for details.

## Discussion

Anticipating a growing knowledge-base of the molecular determinants of antibiotic resistance (1), we have developed a method of extracting and purifying mycobacterial DNA from primary clinical samples and producing accurate sequence data in clinically a useful timeframe. We have demonstrated: first, that direct WGS of sputum is possible, and gives concordant DST results with phenotype and concordant phylogenetic placement with culture-based sequencing. Second: using an Illumina MiSeq sequencer, we can obtain results within 48 hours for <£100 per sample. Using an Illumina MiniSeq can deliver a same-day test (16 hours) for an estimated consumable cost of £198 per sample. Although the costs presented here only represent reagents, they are well below that of traditional phenotyping (£545 to provide first-/second- line DST and MIRU-VNTR in a bottom-up costing including, for example, consumables, staff time, and overheads (16)). The MiSeq cost is also well below that of the SureSelect procedure (£203 per sample) (18).

The World Health Organisation (WHO) has called for affordable and accessible point-of-care TB diagnostics, including for DST. Current molecular assays provide partial information on some drugs, but do not easily scale to incorporate a growing list of recognized resistance mutations. Furthermore, additional assays are currently needed where surveillance or outbreak detection are indicated, at additional cost. A single assay providing diagnostic information, and data for surveillance and outbreak detection is therefore an attractive prospect.

In cities where there are large numbers of TB cases (for example upward of 65,000 TB cases per years in Mumbai) centralized sequencing services taking advantage of high throughput Illumina sequencing platforms may be applicable. However, the relatively high capital costs, and requirement for a well-equipped laboratory are an impediment to implementation across the full range of locations across the world. For a complete solution, the ability to function in varied low-tech environments is a practical necessity. The MinION delivers this, as demonstrated in Guinea last year during the Ebola outbreak (30). We confirm here that, despite the high error rate in reads, given deep coverage, it is possible to accurately genotype resistance SNPs and indels using the MinION method applied here.

Since with Illumina technology the depth of sequencing is determined in advance, the small amount of *M. tuberculosis* in a direct sample can result in test failures. In this experiment *M. tuberculosis* identification and susceptibility prediction failed in 2/39 and 13/37 samples respectively. MinION sequencing allows sequencing to continue until sufficient coverage has been obtained, giving faster results when there is high load, and avoiding this type of failure when the load is low. The throughput obtained here with R9.4 flowcells (1.3 Gb), would allow an 8 hour turnaround time with one sample per flowcell. Overall yield gave BCG coverage depth of 147x, sufficient to have multiplexed 3-5 samples, with implied cost of around £101-172/sample - more data is needed to confidently determine a realistic multiplexing level. Although these estimated costs are not yet at a level to be affordable in routine global use, there is clearly scope for further technology-driven cost reduction, either through improved enrichment/depletion (Figure 2a), higher sequencing yield and multiplexing, or real-time filtering of contamination (31).

Direct from patient sequencing may also allow identification of mycobacterial co-colonization, which may occur in 3-5% of patients (32). Indeed, since infections are chronic and structured in the lung, it is expected that transmission models will need to account for mixtures and within-host evolution (33).

Diagnostic and surveillance information direct from patient specimens can now be obtained in as little as 8 hours on the ONT sequencing platform, and in 16/44 hours on Illumina platforms, a considerable step forward. Faster and more automated sample processing, as well as a cost reduction, is a clear necessity for global take-up in low-income settings. Achieving this would revolutionise the management of TB.

## Acknowledgements

We thank Phuong Quan for assistance with statistical analysis, Rachel Norris for help with error analysis, Kevin Hall and Aurelie Modat from Illumina for helping with the MiniSeq sequencing, and David Stoddart and Oliver Hartwell from Oxford Nanopore Technologies for giving us help and early access to the rapid PCR 1D prep.

## References

1. Walker TM, Kohl TA, Omar SV, Hedge J, del Ojo Elias C, Bradley P, Iqbal Z, Feuerriegel S, Niehaus KE, Wilson DJ, Clifton DA, Kapatai G, Ip CLC, Bowden R, Drobniewski FA, Allix-Beguec C, Gaudin C, Parkhill J, Diel R, Supply P, Crook DW, Smith EG, Walker AS, Ismail N, Niemann S, Peto TEA, Modernising Medical Microbiology (MMM) Informatics Group (2015) Whole-genome sequencing for prediction of Mycobacterium tuberculosis drug susceptibility and resistance: a retrospective cohort study. The Lancet Infectious Diseases 15:1193–202

2. Said HM, Kock MM, Ismail NA, Baba K, Omar SV, Osman AG, Hoosen AA, Ehlers MM (2012) Evaluation of the GenoType MTBDRsl assay for susceptibility testing of second-line anti-tuberculosis drugs. Int J Tuberc Lung Dis. 2012 Jan;16(1):104–9

3. WHO Expert Group Report (2008) Molecular Line Probe Assays for Rapid Screening of Patients at Risk of Multi-Drug Resistant Tuberculosis (MDR-TB)

4. Colman RE, Anderson J, Lemmer D, Lehmkuhl E, Georghiou SB, Heaton H, Wiggins K, Gillece JD, Schupp JM, Catanzaro DG, Crudu V, Cohen T, Rodwell TC, Engelthaler DM (2016) Rapid drug susceptibility testing of drug-resistant *Mycobacterium tuberculosis* isolates directly from clinical samples by use of amplicon sequencing: a concept study. J Clin Microbiol 54:2058–2067

5. Lee RS, Behr MA (2016) The implications of whole-genome sequencing in the control of tuberculosis. Therapeutic advances in infectious disease 3:47–62

6. Takiff HE, Feo O (2015) Clinical value of whole-genome sequencing of Mycobacterium tuberculosis. Lancet Infect Dis 15:1077–90

7. Witney AA, Cosgrove CA, Arnold A, Hinds J, Stoker NG, Butcher PD (2016) Clinical use of whole genome sequencing for Mycobacterium tuberculosis. BMC medicine 14:46

8. Casali N, Nikolayevskyy V, Balabanova Y, Harris SR, Ignatyeva O, Konsevaya I, Corander J, Bryant J, Parkhill J, Nejentsev S, Horstmann RD, Brown T, Drobniewski F (2014) Evolution and transmission of drug-resistant tuberculosis in a Russian population. Nat Genet 46:279–86

9. Clark TG, Mallard K, Coll F, Preston M, Assefa S, Harris D, Ogwang S, Mumbowa F, Kirenga B, O’Sullivan DM, Okwera A, Eisenach KD, Joloba M, Bentley SD, Ellner JJ, Parkhill J, Jones-Lopez EC, McNerney R (2013) Elucidating emergence and transmission of multidrug-resistant tuberculosis in treatment experienced patients by whole genome sequencing. PLoS One 8:e83012

10. Farhat MR, Shapiro BJ, Kieser KJ, Sultana R, Jacobson KR, Victor TC, Warren RM, Streicher EM, Calver A, Sloutsky A, Kaur D, Posey JE, Plikaytis B, Oggioni MR, Gardy JL, Johnston JC, Rodrigues M, Tang PKC, Kato-Maeda M, Borowsky ML, Muddukrishna B, Kreiswirth BN, Kurepina N, Galagan J, Gagneux S, Birren B, Rubin EJ, Lander ES, Sabeti PC, Murray M (2013) Genomic analysis identifies targets of convergent positive selection in drug-resistant Mycobacterium tuberculosis. Nat Genet 45:1183–9

11. Gardy JL, Johnston JC, Ho Sui SJ, Cook VJ, Shah L, Brodkin E, Rempel S, Moore R, Zhao Y, Holt R, Varhol R, Birol I, Lem M, Sharma MK, Elwood K, Jones SJM, Brinkman FSL, Brunham RC, Tang P (2011) Whole-genome sequencing and social-network analysis of a tuberculosis outbreak. The New England journal of medicine 364:730–9

12. Guerra-Assuncao JA, Crampin AC, Houben RM, Mzembe T, Mallard K, Coll F, Khan P, Banda L, Chiwaya A, Pereira RP, McNerney R, Fine PE, Parkhill J, Clark TG, Glynn JR (2015) Large-scale whole genome sequencing of M. tuberculosis provides insights into transmission in a high prevalence area. Elife. 2015 Mar 3;4.

13. Stucki D, Ballif M, Bodmer T, Coscolla M, Maurer AM, Droz S, Butz C, Borrell S, Langle C, Feldmann J, Furrer H, Mordasini C, Helbling P, Rieder HL, Egger M, Gagneux S, Fenner L (2015) Tracking a tuberculosis outbreak over 21 years: strain-specific single-nucleotide polymorphism typing combined with targeted whole-genome sequencing. J Infect Dis 211:1306–16

14. Walker TM, Ip CL, Harrell RH, Evans JT, Kapatai G, Dedicoat MJ, Eyre DW, Wilson DW, Hawkey PM, Crook DW, Parkhill J, Harris D, Walker AS, Bowden R, Monk P, Smith EG, Peto TE (2013) Whole-genome sequencing to delineate Mycobacterium tuberculosis outbreaks: a retrospective observational study. Lancet Infect Dis 13:137–46

15. Walker TM, Lalor MK, Broda A, Saldana Ortega L, Morgan M, Parker L, Churchill S, Bennett K, Golubchik T, Giess AP, Del Ojo Elias C, Jeffery KJ, Bowler IC, Laurenson IF, Barrett A, Drobniewski F, McCarthy ND, Anderson LF, Abubakar I, Thomas HL, Monk P, Smith EG, Walker AS, Crook DW, Peto TE, Conlon CP (2014) Assessment of Mycobacterium tuberculosis transmission in Oxfordshire, UK, 2007-12, with whole pathogen genome sequences: an observational study. Lancet Respir Med 2014 Apr;2(4):285–92

16. Pankhurst LJ, Del Ojo Elias C, Votintseva AA, Walker TM, Cole K, Davies J, Fermont JM, Gascoyne-Binzi DM, Kohl TA, Kong C, Lemaitre N, Niemann S, Paul J, Rogers TR, Roycroft E, Smith EG, Supply P, Tang P, Wilcox MH, Wordsworth S, Wyllie D, Xu L, Crook DW, COMPASS-TB Study Group (2016) Rapid, comprehensive, and affordable mycobacterial diagnosis with whole-genome sequencing: a prospective study. Lancet Respir Med 2016 Jan;4(1):49–58

17. Doughty EL, Sergeant MJ, Adetifa I, Antonio M, Pallen MJ (2014) Culture-independent detection and characterisation of Mycobacterium tuberculosis and M. africanum in sputum samples using shotgun metagenomics on a benchtop sequencer. PeerJ 2014 Sep 23;2:e585

18. Brown AC, Bryant JM, Einer-Jensen K, Holdstock J, Houniet DT, Chan JZ, Depledge DP, Nikolayevskyy V, Broda A, Stone MJ, Christiansen MT, Williams R, McAndrew MB, Tutill H, Brown J, Melzer M, Rosmarin C, McHugh TD, Shorten RJ, Drobniewski F, Speight G, Breuer J (2015) Rapid Whole-Genome Sequencing of Mycobacterium tuberculosis Isolates Directly from Clinical Samples. J Clin Microbiol 53:2230–7

19. Votintseva AA, Pankhurst LJ, Anson LW, Morgan MR, Gascoyne-Binzi D, Walker TM, Quan TP, Wyllie DH, Del Ojo Elias C, Wilcox M, Walker AS, Peto TE, Crook DW (2015) Mycobacterial DNA extraction for whole-genome sequencing from early positive liquid (MGIT) cultures. J Clin Microbiol 53:1137–43

20. Li, H. (2013) Aligning sequence reads, clone sequences and assembly contigs with BWA-MEM. arXiv:1303.3997

21. Bradley P, Gordon NC, Walker TM, Dunn L, Heys S, Huang B, Earle S, Pankhurst LJ, Anson L, de Cesare M, Piazza P, Votintseva AA, Golubchik T, Wilson DJ, Wyllie DH, Diel R, Niemann S, Feuerriegel S, Kohl TA, Ismail N, Omar SV, Smith EG, Buck D, McVean G, Walker AS, Peto TE, Crook DW, Iqbal Z (2015) Rapid antibiotic-resistance predictions from genome sequence data for Staphylococcus aureus and Mycobacterium tuberculosis. Nat. Commun. 2015 Dec 21;6:10063

22. Iqbal Z, Caccamo M, Turner I, Flicek P, McVean G (2012) De novo assembly and genotyping of variants using colored de Bruijn graphs. Nat Genet 2012 Jan 8; 44(2):226–232

23. Loman NJ, Quinlan AR (2014) Poretools: a toolkit for analysing nanopore sequence data. Bioinformatics 2014 Dec 1;30(23):3399–401

24. Vaser R, Sovic I, Nagarajan N, Sikic M (2016) Fast and accurate de novo assembly from long uncorrected reads bioRxiv http://dx.doi.org/10.1101/068122

25. Kurtz S, Phillippy A, Delcher AL, Smoot M, Shumway M, Antonescu C, Salzberg S (2004) Versatile and open software for comparing genomes

26. Koren S, Walenz BP, Berlin K, Miller JR, Phillippy AM (2016) Canu: scalable and accurate long read assembly via adaptive k-mer weighting and repeat separation. bioRxiv: http://dx.doi.org/10.1101/071282

27. Cohen KA, Abeel T, Manson McGuire A, et al. (2015) Evolution of Extensively Drug-Resistant Tuberculosis over Four Decades: Whole Genome Sequencing and Dating Analysis of Mycobacterium tuberculosis Isolates from KwaZulu-Natal. PLoS medicine 12:e1001880

28. Sanchez-Padilla E, Merker M, Beckert P, et al. (2015) Detection of drugresistant tuberculosis by Xpert MTB/RIF in Swaziland. N Engl J Med 2015 Mar 19;372(12):1181–2

29. Eldholm V, Norheim G, Lippe Bvd, Kinander W, Dahle UR, Caugant DA, Mannsaker T, Mengshoel AT, Dyrhol-Riise AM, Balloux F (2014) Evolution of extensively drug-resistant Mycobacterium tuberculosis from a susceptible ancestor in a single patient. Genome Biol. 2014; 15(11): 490.

30. Quick J, Loman NJ, Duraffour S, Simpson JT, Severi E, Cowley L, Bore JA, Koundouno R, Dudas G, Mikhail A, Ouédraogo N, Afrough B, Bah A, Baum JH, Becker-Ziaja B, Boettcher JP, Cabeza-Cabrerizo M, Camino-Sánchez Á, Carter LL, Doerrbecker J, Enkirch T, García-Dorival I, Hetzelt N, Hinzmann J, Holm T, Kafetzopoulou LE, Koropogui M, Kosgey A, Kuisma E, Logue CH, Mazzarelli A, Meisel S, Mertens M, Michel J, Ngabo D, Nitzsche K, Pallasch E, Patrono LV, Portmann J, Repits JG, Rickett NY, Sachse A, Singethan K, Vitoriano I, Yemanaberhan RL, Zekeng EG, Racine T, Bello A, Sall AA, Faye O, Faye O, Magassouba N, Williams CV, Amburgey V, Winona L, Davis E, Gerlach J, Washington F, Monteil V, Jourdain M, Bererd M, Camara A, Somlare H, Camara A, Gerard M, Bado G, Baillet B, Delaune D, Nebie KY, Diarra A, Savane Y, Pallawo RB, Gutierrez GJ, Milhano N, Roger I, Williams CJ, Yattara F, Lewandowski K, Taylor J, Rachwal P, Turner DJ, Pollakis G, Hiscox JA, Matthews DA, O’Shea MK, Johnston AM, Wilson D, Hutley E, Smit E, Di Caro A, Wölfel R, Stoecker K, Fleischmann E, Gabriel M, Weller SA, Koivogui L, Diallo B, Keïta S, Rambaut A, Formenty P, Günther S, Carroll MW (2016) Real-time, portable genome sequencing for Ebola surveillance. Nature 2016 Feb 11;530(7589):228–32

31. Loose M, Malla S, Stout M (2016): Real-time selective sequencing using nanopore technology. Nat Methods 2016 Sep;13(9):751–4

32. Wyllie D, Bawa Z, Walker T, Peto T, Smoth G (2016): Low frequency of mixed M. tuberculosis infection in a large prospective UK series as assessed using whole genome sequencing. 26^th^ ECCMID Amsterdam, Netherlands, 9–12 April 2016 P0153

33. Lieberman T, Wilson D, Misra R, Xiong LL, Moodley P, Cohen T, Kishony R (2016) Genomic diversity in autopsy samples reveals within-host dissemination of HIV-associated Mycobacterium tuberculosis. Nat Med 2016 Oct 31

